# Novel anionic cecropins from the spruce budworm feature a poly-L-aspartic acid C-terminus

**DOI:** 10.1101/307702

**Authors:** Halim Maaroufi, Michel Cusson, Roger C. Levesque

**Affiliations:** Institut de biologie intégrative et des systèmes (IBIS), Université Laval, Quebec City, Canada; Natural Resources Canada, Canadian Forest Service, Laurentian Forestry Centre, Quebec City, Canada; Institut de biologie intégrative et des systèmes (IBIS) and Faculté de médecine, Université Laval, Quebec City, Canada

**Keywords:** *Choristoneura fumiferana*, anionic cecropins, C-terminal poly-L-aspartic acid, ancient duplication, apoptotic motif, anticancer peptide

## Abstract

Cecropins form a family of amphipathic α-helical cationic peptides with broad-spectrum antibacterial properties and potent anticancer activity. The emergence of bacteria and cancer cells showing resistance to cationic antimicrobial peptides (CAMPs) has fostered a search for new, more selective and more effective alternatives to CAMPs. With this goal in mind, we looked for cecropin homologs in the genome and transcriptome of the spruce budworm, *Choristoneura fumiferana*. Not only did we find paralogs of the conventional cationic cecropins (Cfcec^+^), our screening also led to the identification of previously uncharacterized anionic cecropins (Cfcec^−^), featuring a poly-L-aspartic acid C-terminus. Comparative peptide analysis indicated that the C-terminal helix of Cfcec^−^ is amphipathic, unlike that of Cfcec^+^, which is hydrophobic. Interestingly, molecular dynamics simulations pointed to the lower conformational flexibility of Cfcec^−^ peptides, relative to that of Cfcec^+^. Phylogenetic analysis suggests that the evolution of distinct Cfcec^+^ and Cfcec^−^ peptides may have resulted from an ancient duplication event within the Lepidoptera. Our analyses also indicated that Cfcec^−^ shares characteristics with entericidins, which are involved in bacterial programmed cell death, lunasin, a peptide of plant origins with antimitotic effects, and APC15, a subunit of the anaphase-promoting complex. Finally, we found that both anionic and cationic cecropins contain a BH3-like motif (G-[KQR]-[HKQNR]-[IV]-[KQR]) that could interact with Bcl-2, a protein involved in apoptosis; this observation is congruent with previous reports indicating that cecropins induce apoptosis. Altogether, our observations suggest that cecropins may provide templates for the development of new anticancer drugs.

**Graphical abstract:** 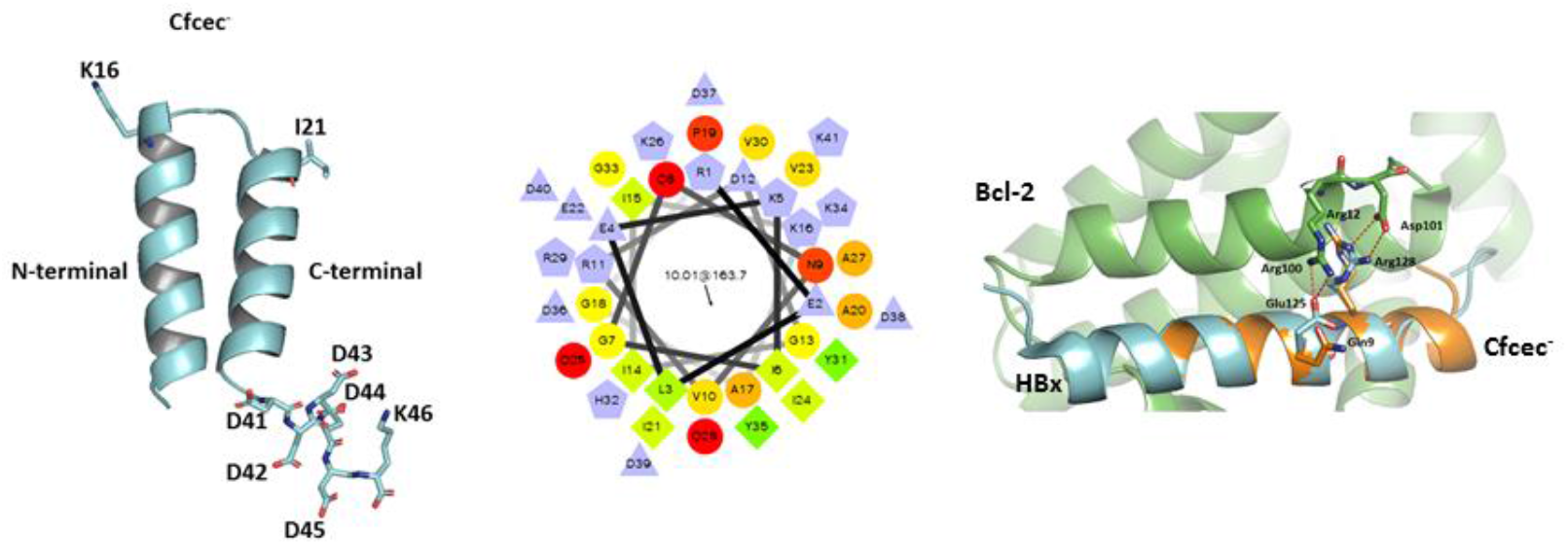

**Highlights:**

1. Genes encoding novel anionic cecropins (Cfcec^−^), featuring a C-terminal poly-L-aspartic acid, were found in the genome of the spruce budworm, *Choristoneura fumiferana*.
2. Divergence between Cfcec^+^ and Cfcec^−^ could be the result of an ancient duplication event within the Lepidoptera.
3. There is an apparent relationship between motifs observed in cecropin peptides and apoptosis.
4. Anionic cecropins from the spruce budworm display characteristics suggesting they could have anticancer activity

## 1. Introduction

Antimicrobial peptides (AMPs) constitute an important component of the innate immune system of insects and, as such, may have played a role in the evolutionary success of this highly speciose taxon, whose members occupy almost all habitats in nature. To date, over 150 insect AMPs have been identified (Yi et al., 2014), with cecropin being the first one to have been purified, in 1980, from pupae of the cecropia moth, *Hyalophora cecropia* (Hultmark et al., 1980; Steiner et al., 1981). Cecropins are now known to form a family of amphipathic α-helical peptides containing 34–39 amino acid residues. They display a broad spectrum of antibacterial properties and act as modulator of the innate immune system. They also show potent anticancer activity (Suttmann et al., 2008; Hoskin and Ramamoorthy, 2008; Huang et al., 2015).

Cecropins characterized to date fall in the category of cationic antimicrobial peptides (CAMPs), whose distinguishing feature is an excess in basic amino acids (positive charge, cationic; Otvos, 2000). Their amphipathic properties allow interactions with membranes (Lee et al., 2013) and their toxicity toward bacteria is considered to be primarily due to an initial electrostatic interaction between the peptide and the anionic phospholipid head groups in the outer layer of the bacterial cytoplasmic membrane, ultimately causing membrane disruption (Zasloff, 2002).

For the present work, we took advantage of genomic and transcriptomic resources recently developed by our group for the spruce budworm, *Choristoneura fumiferana,* to identify and characterize the repertoire of cecropins in this important lepidopteran conifer pest. Our analyses led to the identification of novel, anionic cecropins (Cfcec^−^) featuring a poly-L-aspartic acid (PAA) C-terminus. We used *in silico* approaches to compare the physico-chemical properties of the anionic peptides with those of the more conventional cationic cecropins, some of which were also found in the *C. fumiferana* genome (Cfcec^+^). In addition, we compared *C. fumiferana* cecropins with other bioactive peptides whose biochemical properties have been well characterized; in the process, we identified some peptides that share features with cecropins, including motifs that could explain why the latter can induce apoptosis.

## 2. Materials and Methods

### 2.1. C. fumiferana genomic and transcriptomic resources

To identify spruce budworm cecropins, we queried a draft assembly of the *C. fumiferana* genome, generated from Roche 454 (GS-FLX+) whole genome shotgun reads, assembled using the Newbler software (Roche); sequencing was performed using DNA extracted from a single male pupa (Cusson et al. unpublished). Similar searches were conducted by querying a *C. fumiferana* transcriptome. The latter was generated using both Illumina and Roche 454 (GS-FLX+) reads following sequencing of a normalized cDNA library, generated from a pool of mRNAs collected from all *C. fumiferana* life stages. Contig assembly was carried out using the MIRA (Chevreux et al. 2004) assembler (Brandão et al. unpublished).

### 2.2. Blast

Similarity searches in our genomic and transcriptomic databases were performed locally using the tblastn algorithm (http://www.ncbi.nlm.-nih.gov/blast) and the sequence of the *H. cecropia* cecropin-A (HccecA; Uniprot: P01507) as query. We also conducted similar tblastn searches against a public *C. fumiferana* EST database (NCBI). To determine if other cecropin homolog sequences with PAA exist in other species, we searched in all sequenced genomes in GenBank using BLASTp, tBLASTn and HMM profiles.

### 2.3. Chemical properties of cecropins

The chemical structures and properties of cecropin peptides were investigated *in silico* using PepDraw (http://www.tulane.edu/~biochem/WW/PepDraw/index.html). Hydrophobicity, as determined by PepDraw, is the free energy associated with transitioning a peptide from an aqueous environment to a hydrophobic environment such as octanol. The scale used is the Wimley-White scale, an experimentally determined scale, where the hydrophobicity of the peptide is the sum of Wimley-White hydrophobicities and measured in Kcal/mol (White and Wimley, 1998). Neutral pH is assumed.

### 2.4. Structure predictions and molecular dynamics simulations

The secondary structures of cecropins and lunasin were predicted using Psipred software (Jones, 1999), while helical wheel projections for the same peptides were performed using the tool available at http://rzlab.ucr.edu/scripts/wheel/wheel.cgi.

3D homology models of *C. fumiferana* cecropins and lunasin were built using the crystal structures of papiliocin of *Papilio xuthus* (PDB id: 2LA2) and allergen Arah6 of *Arachis hypogaea* (PDB id: 1W2Q) as templates, respectively, using the modeling software Modeller (Webb and Sali, 2014). Model quality was assessed by Ramachandran plot analysis through PROCHECK (Laskowski et al., 1993). Structure images were generated using PyMOL (http://www.pymol.org).

In order to evaluate conformational changes of cecropins, molecular dynamics (MD) simulations were performed in GROMACS (v5.1.4) using the OPLS-AA/L all-atom force field (Kaminski et al. 2001). Cecropins were solvated in a cubic box as the unit cell, using SPC/E water model with the box edge distance from the molecule set to 1.0 nm. The system was neutralized by replacing solvent molecules with Cl^−^ and Na^+^ ions. Energy minimization was conducted using the steepest descent method to ensure that the system has no steric clashes or inappropriate geometry. Equilibration of the solvent and ions around the peptide was conducted under NVT (300 K) and NPT (1.0 bar) ensembles for 100 ps. Cecropin MD simulations were conducted for 1 ns.

### 2.5. Phylogenetic analysis

We used the amino acid sequences of cationic and anionic cecropins to search by BlastP for close homologs in insects. To assess phylogenetic relationships among cecropins of *C. fumiferana* and those of Diptera, Coleoptera and other Lepidoptera, sequences were aligned using Muscle (Edgar, 2004) and a phylogenetic tree was constructed using the Neighbor-Joining method (Saitou and Nei, 1987). Evolutionary distances were computed using the Equal Input method (Tajima and Nei, 1984) and are shown as the number of amino acid substitutions per site. All positions displaying less than 95% site coverage were eliminated. Phylogenetic analyses were conducted in MEGA6 (Tamura et al., 2013).

## 3. Results and discussion

### 3.1. *Search for* C. fumiferana *cecropins in genomic and transcriptomic resources*

To identify cecropin orthologs in the sequenced genome and transcriptome of *C. fumiferana* (unpublished data), we conducted tblastn searches using cecropin-A of *H. cecropina* (Uniprot: P01507) as query. We found two types of cecropin genes that were designated *Cfcec*^+^ (cationic cecropins) and *Cfcec*^−^ (anionic cecropins). Cfcec^+^ and Cfcec^−^ peptides were again used as queries to carry out tblastn iteratively until no new hit occurred. Two *Cfcec*^+^ and two *Cfcec*^−^ genes were found in the genome and transcriptome of *C. fumiferana*. These genes were designated *Cfcec*^+^1, *Cfcec*^+^2 and *Cfcec*^−^1, *Cfcec*^−^2, for cationic and anionic cecropins, respectively (Table 1; Fig. 1). Cecropin genes are composed of two exons and one intron (Fig. 1A, B). In the draft assembly of the *C. fumiferana* genome, the *Cfcec*^+^1 and *Cfcec*^−^1 genes were localized on the same scaffold with an interval of 3588 bp, whereas the *Cfcec*^+^2 and *Cfcec*^−^2 genes were found on distinct scaffolds. Interestingly, another scaffold contains the exon 1 of *Cfcec*^+^2 and the C-terminal poly-L-aspartic acid (PAA) of *Cfcec*^−^1 (Fig. 1B). It is tempting to speculate that the ancestor of *Cfcec*^−^ acquired the PAA C-terminus through exon shuffling.

**Tableau 1.**
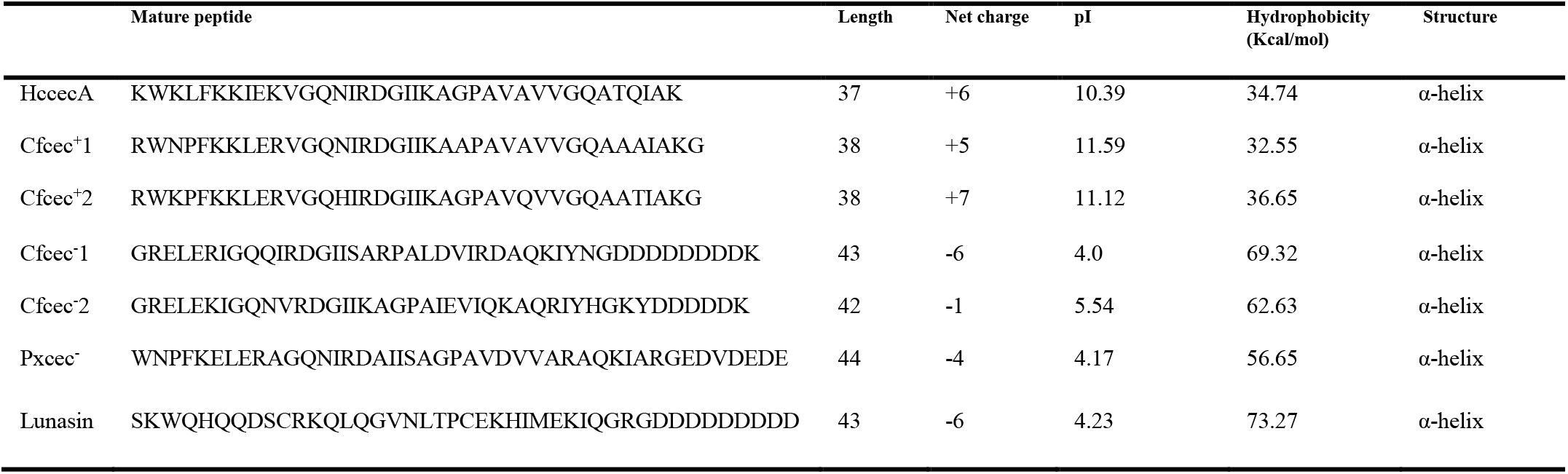
Physico-chemical characteristics of HccecA, Cfcec^+^, Cfcec^−^, Pxcec^−^ and lunasin.

**Figure 1.**
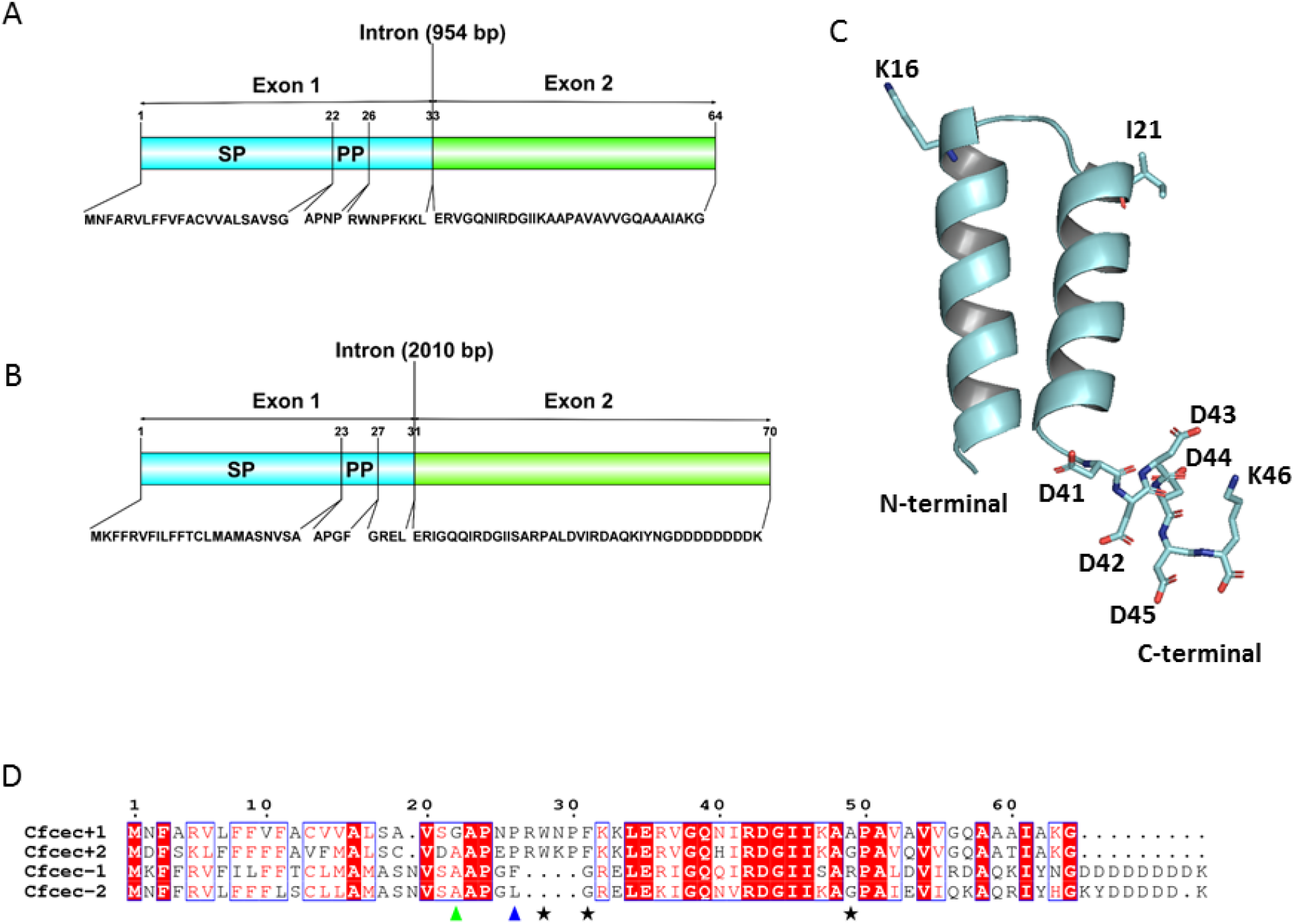
(A) Comparison of *Cfcec*^+^1 and (B) *Cfcec*^−^1genes. The genes are composed of two exons and one intron. Amino acids sequence of prepeptides are indicated below each gene. SP, signal peptide; PP, propeptide. The figure was generated with DOG 1.0 (Ren et al. 2009). (C) 3D model of anionic cecropin Cfcec^−^2. (D) Alignment of Cfcec^+^ and Cfcec^−^ peptides. Green and blue arrows identify the C-terminal residue of the signal peptide (SP) and the propeptide (PP), respectively. Black stars represent functionally important amino acids. Muscle (Edgar 2004) was used to create multiple alignments. The figure was prepared with ESPript (http://espript.ibcp.fr).

To confirm that *Cfcec*^+^ and *Cfcec*^−^ are actually transcribed, we searched *C. fumiferana* transcriptomic and EST databases, where we found the corresponding transcripts. In addition, NCBI”s EST database revealed that Cfcec^+^ and some anionic cecropins of Lepidoptera (Fig. S1) were expressed in frontline defense tissues such as the fat body, epidermis and midgut, as shown for *Plutella xylostella* (Jin et al., 2012).

### 3.2. Peptide analysis

#### 3.2.1. Sequence analysis

Alignment of Cfcec^+^ and Cfcec^−^ showed that the major differences between these two peptides are found at the N- and C-termini of the mature peptides (Fig. 1D). Cfcec^+^2 displays characteristics similar to those of other cecropins of moths where the first residue, preceding the conserved Trp2, is a Lys or an Arg (Otvos, 2000). This is not the case for Cfcec^−^ where these amino acid residues are absent (Fig. 1D). The Trp2 residue of HccecA was shown to be important for activity against all tested bacteria (Andreu et al., 1985). Trp2 and Phe5 of *Papilio xuthus* cecropin (Pxcec^+^) were also shown to interact with LPS (Kim et al., 2011). These two amino acid residues are conserved in Cfcec^+^ but not in Cfcec^−^ (Fig.1D). In addition, the hinge region of cecropins (Gly-Pro) is conserved in Cfcec^+^2 and Cfcec^−^2 as described by Efimova et al. (2014) and provides conformational flexibility (Oh et al., 2000) (Fig.1D). Analysis of amino acid residues of both types of *C. fumiferana* cecropins suggests that Cfcec^+^ shares many characteristics with HccecA (Table 1).

Blast searches were conducted to determine whether anionic cecropins such as Cfcec^−^ were present in other insects. Not surprisingly, we found a few cecropins bearing three or four acidic amino acid residues at their C-termini, conferring a negative net charge (−2 to −4) to the peptides (Fig. S1). The characteristics of anionic *P. xuthus* cecropin (Pxcec^−^) are presented in Table 1; this peptide shares several characteristics with Cfcec^−^, including a similar length, a negative net charge, and comparable pI and hydrophobicity values. To our knowledge, the activity and structure of these anionic cecropins has never been examined experimentally. Finally, it is worth noting that the PAA of Cfcec^−^ ends with a lysine residue (Fig. 2). Interestingly, a lysine residue was similarly found at the end of the poly-L-aspartic acid peptides (VDDDDK, APDDDDK and TDDDK) studied by Brogden et al. (1997). It is conceivable that the role of this positively charged residue is to interact with negative charges on membrane. Indeed, the long nonpolar region of the side chain of lysine was shown to extend or snorkel into the hydrophobic core of the target membrane (Li et al., 2013).

**Figure 2.**
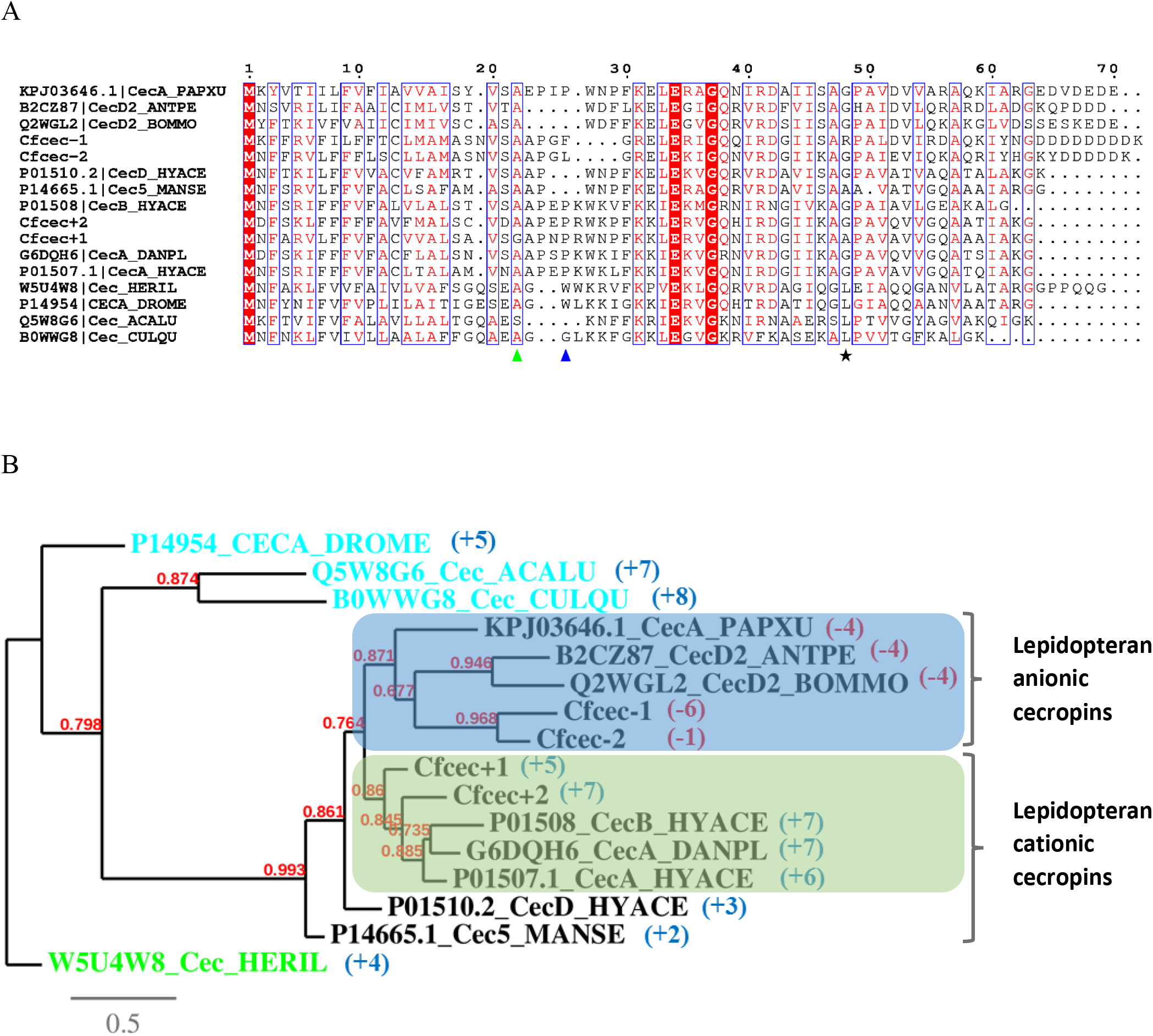
(A) Multiple sequence alignment and (B) phylogenetic relationship among cecropins (based on amino acid sequences). Green letters: Coleoptera (represented by *Acalolepta luxuriosa* (ACALU); blue letters: Diptera (represented by *Drosophila melanogaster* (DROME), *Hermetia illucens* (HERIL) and *Culex quinquefasciatus* (CULQU). Black letters: Lepidoptera (represented by *Antheraea pernyi* (ANTPE), *Bombyx mori* (BOMMO), *Danaus plexippus* (DANPL), *Hyalophora cecropia* (HYACE) *Manduca sexta* (MANSE), and the *C. fumiferana* cecropins identified in this work. Net charge of each mature peptide is in brackets. The distinct anionic and cationic cecropin clades are highlighted. The evolutionary history was inferred using the Neighbor-Joining method. The percentage of replicate trees in which the associated cecropins clustered together in the bootstrap test (1000 replicates) are shown next to the branches. The tree was rooted using a coleopteran cecropin as outgroup. Evolutionary analyses were conducted in MEGA6 (Tamura et al. 2013).The multiple alignment figure was prepared with ESPript (http://espript.ibcp.fr).

#### 3.2.2. Structure and molecular dynamics simulation

2D and 3D structures of Cfcec^+^ and Cfcec^−^ were estimated using Psipred and homology modeling (Fig. 1C), respectively. For both types of peptides, the structure comprises two helices, as shown by NMR for other cecropins (PDB id: 2LA2 and 2MMM). To illustrate the properties of α-helices in these peptides, we constructed helical wheel diagrams. Thus, the N-terminal helices of Cfcec^+^ and Cfcec^−^ both display amphipathic properties (amino acids 1-21 of Cfcec^+^ and 1-17 of Cfcec^−^), whereas their C-terminal helices (amino acids 25–37 of Cfcec^+^ and 21-34 of Cfcec^−^) are hydrophobic and amphipathic, respectively (Fig. S2). In comparison, lunasin, a peptide of plant origins that shares some characteristics with Cfcec^−^ (Table 1), has N- and C-terminal helices that are hydrophilic and amphipathic, respectively (Fig. S2). The high antibacterial activity of cecropin-like model peptides has been shown to require a basic, amphipathic N-terminal helix and a hydrophobic C-terminal helix, connected by a flexible hinge region (Fink et al., 1989). Cfcec^+^ possesses these characteristics and can therefore be classified as a CAMP. However, Cfcec^−^ does not fit this pattern, with its amphipathic C-terminal helix, a feature shared with lunasin (Dia and de Mejia, 2011), which displays anti-cancer activity.

Molecular dynamics simulations showed that Cfcec^−^ peptides have less conformational flexibility at their N- and C-termini than Cfcec^+^ (Fig. S3), with only the PAA of Cfcec^−^ displaying significant flexibility. Flexibility appears to be provided primarily by glycine residues (Fig. S3). Since CAMP activity is dependent upon the presence of conformational flexibility (Amos et al. 2016), differences in this variable between the Cfcec^+^ and Cfcec^−^ peptides, in addition to the different physico-chemical properties of their C-termini, suggests that Cfcec^+^ and Cfcec^−^ could have different biological activities.

### 3.3. Phylogeny

To infer phylogenetic relationships among cecropins of *C. fumiferana* and those of other Lepidoptera, Diptera, and Coleoptera, sequences reported here and others gleaned from public databases were aligned using Muscle (Edgar, 2004), and a phylogenetic tree was constructed using the Neighbor-Joining method (Saitou and Nei, 1987). The branching pattern obtained suggests that Cfcec^+^ peptides are more closely related to their *H. cecropia* and *D. plexipus* CecA and CecB orthologs than to their *C. fumiferana* paralogs (Cfcec^−^; Fig. 2B). Again, this observation suggests that anionic cecropins could have arisen following an ancient duplication event within the Lepidoptera. Not surprisingly, lepidopteran cecropins with negative and positive net charges formed two separate clades (Figure 2B). It has been observed that the net charge of α-helical ACPs has an effect on the anticancer activity of the peptide, with an increase in the net charge enhancing anticancer activity (Huang et al., 2015).

### 3.4. Peptides that display similarity to Cfcec^+^ and Cfcec^−^

In recent years, AMPs have become promising molecules to fight cancer (Hoskin and Ramamoorthy, 2008; Riedl et al., 2011). The Antimicrobial Peptide Database (APD, http://aps.unmc.edu/AP/main.php) currently contains 193 peptides that are identified as anticancer-peptides (ACPs). These peptides, however, are from different sources and display limited similarity to one another. Nonetheless, they share some characteristics, including a positive charge and an amphipathic helix, and they exhibit a large spectrum of anticancer activity (Gaspar et al., 2013; Lu et al., 2016).

BLAST searches against the AMP database using Cfcec^−^ as query revealed that the PAA of mature lunasin (amino acids 56-64) is similar to that of Cfcec^−^. Moreover, lunasin shares other features with Cfcec^−^, including net charge, pI, hydrophobicity and secondary structure (Table 1). Lunasin has antimitotic properties attributed to the binding of its PAA C-terminus to regions of hypoacetylated chromatin (Galvez and de Lumen, 1999; Galvez et al., 2001). In the AMP database, we also found other short peptides containing mostly aspartic acid residues, including the peptide DEDDD, which shows inhibitory activity against breast cancer cells (Li et al., 2016), as well as three other AMPs: DDDDDDD, GDDDDDD and GADDDDD (Brogden et al., 1996). To refine our search for peptides displaying amino acid patterns similar to those of Cfcec^−^, we turned our attention to profile-profile alignment methods (Xu et al., 2014), which are more accurate and more sensitive than sequence-sequence and sequence-profile alignment methods. In this way, we identified HBx, a hepatitis B virus protein whose C-terminus is similar to the N-terminus of Cfcec^−^ (Fig. 3A). HBx contains a BH3-like motif that folds to form an amphipathic α-helix that binds to the conserved BH3-binding groove of Bcl-2, an anti-apoptotic protein (Ma et al., 2008; Jiang et al., 2016). The BH3-like motif may inhibit the anti-apoptotic action of Bcl-2 through interactions with its conserved BH3-binding groove. Glu125 and Arg128 of HBx (Fig. 3C) each makes a pair of charge-stabilized hydrogen bonds to residues Arg100 and Asp101 in Bcl-2 (Jiang et al., 2016).

**Figure 3.**
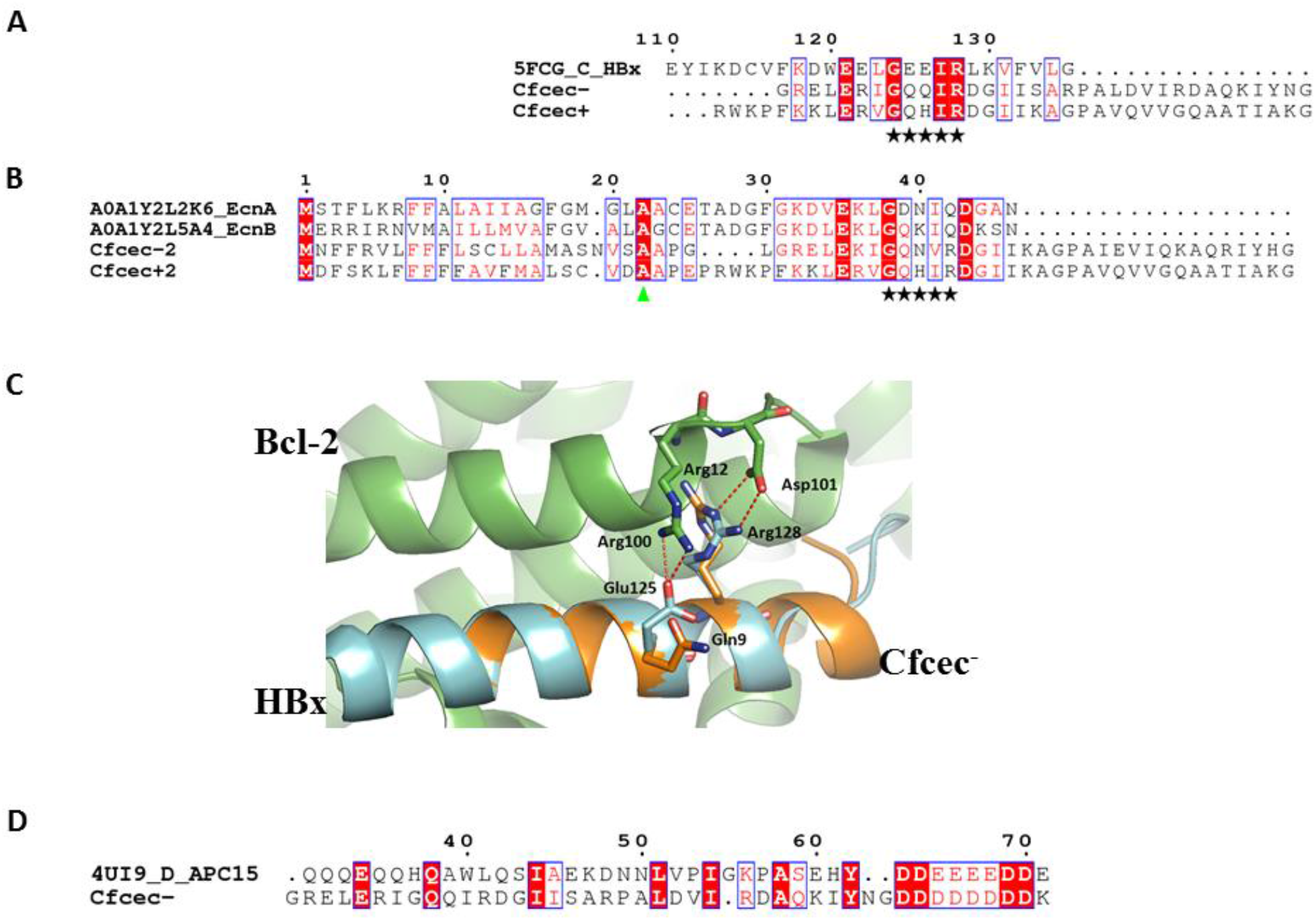
HBx (hepatitis B virus protein), entericidins and APC15 (subunit anaphase-promoting complex 15) sequences display similarities to Cfcec peptides. (A) Alignment of Cfcec^+^ and Cfcec^−^ with HBx. The HBx-like motif is indicated by black stars. (B) Alignment of Cfcec^+^ and Cfcec^−^ with entericidin A and B of *Thalassospira mesophila* (Uniprot: A0A1Y2L2K6 and A0A1Y2L5A4). Green arrow identifies the C-terminal residue of the signal peptide (SP). (C) A close-up of the superposition of 3D models of the N-terminus of Cfcec^−^ and the complex HBx-Bcl-2 (PDBid: 5FCG). N-termini of Cfcec^−^, HBx and Bcl-2 are colored in orange, cyan and green, respectively. (D) Alignment of Cfcec^−^ and APC15. Highly conserved amino acid residues are shown in red and boxed in blue. The sequence alignments and the image of 3D model were prepared with ESPript (http://espript.ibcp.fr) and PyMol (www.pymol.org), respectively.

The amino acids corresponding to Glu125 and Arg128 in the N-terminus of Cfcec^−^ are Gln9 and Arg12 (Fig. 3A and C). To determine if Cfcec^−^ could interact with Bcl-2, we superimposed the 3D model of Cfcec^−^1 onto the crystal structure of the HBx-Bcl-2 complex (PDBid: 5FCG). Indeed, the N-terminus of Cfcec^−^1 superimposed well onto HBx (Fig. 3C) and the amino acids Gln9 and Arg12 could form hydrogen bonds with Arg100 and Asp101 of Bcl-2. Interestingly, a G124L/I127A double mutation (conserved in cecropins; Fig. 3A) in HBx abolished interactions between HBx and CED-9, the functional ortholog of Bcl-2 in *C. elegans* (Geng et al., 2012), pointing to an important functional role of these two residues in HBx, and possibly in cecropins too.

Alignment of cecropins from different species showed that BH3-like motifs are also present at their N-termini. In addition, we found this motif to be present in other AMPs that have been shown to induce apoptosis (Table 2). Interestingly, Choi and Lee (2013) showed that when the first six amino acid residues (GWGSFF) of pleurocidin (Table 2) were truncated, the peptide lost its effect on ROS production (apoptosis), apparently because the N-terminus of pleurocidin contains a BH3-like motif (Table 2, underlined amino acids). Similarly, cecropin-P17 (Table 2) was shown to suppress the proliferation of HepG-2 cells by inducing apoptosis, which was dependent (among other things) on inhibiting Bcl-2 (Wu et al., 2015).

**Tableau 2.**
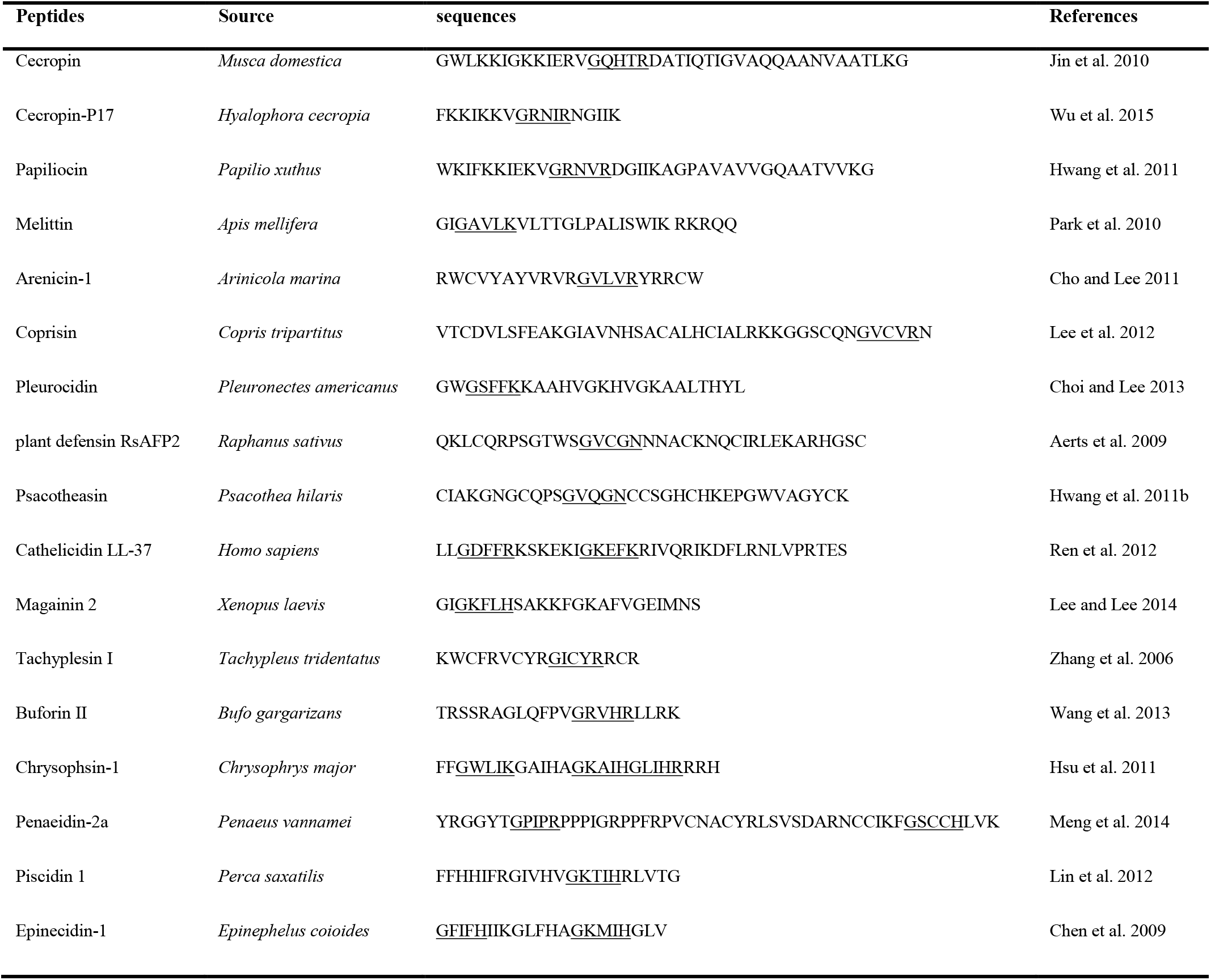
AMPs with shared (G-[KQR]-[HKQNR]-[IV]-[KQR]) motif known to induce apoptosis.

An intensive search for proteins displaying similarity to Cfcec’s led to the finding of entericidins A (EcnA) and B (EcnB) (Fig. 3B), two interesting peptides from bacteria. Entericidins are small lipoproteins encoded by tandem genes whose products function as toxin/antidote in programmed cell death in bacteria. EcnA inhibits the apoptotic action of EcnB by a mechanism that is not yet understood (Bishop et al., 1998). Amino acid sequences of EcnA and EcnB display similarities to AMPs. They have a signal peptide, adopt amphipathic α-helical structures and reciprocally modulate membrane stability (Fig. 3B). Moreover, EcnA and EcnB, like many AMPs, contain a BH3-like motif (Table 2). This supports the hypothesis that programmed cell death genes may have originated in bacteria from a pool of antibiotic genes (Ameisen, 1996).

Lastly, we found that Cfcec^−^ displays sequence similarity to the subunit anaphase-promoting complex 15 (APC15), which plays a role in the release of the mitotic checkpoint complex (MCC) from the APC/C (Fig. 3D). The function of APC15 in human cells seems to be primarily linked to the spindle assembly checkpoint (SAC), and its depletion prevents mitotic slippage (Mansfeld et al., 2011).

### 3.5. Prospective function of PAA

The number of aspartic acid residues differs between the PAA of Cfcec^−^1 and Cfcec^−^2, with five in the latter and eight in the former. This variation changes the net charge of these peptides. It has been reported that variation in the net charge of α-helical ACPs has an effect on the anticancer activity of the peptide (Huang et al., 2015). Moreover, Brogden et al. (1996) showed that the antimicrobial activity of Asp homopolymers increases with the number of Asp residues in the peptide. In addition, lunasin contains a unique Arg-Gly-Asp (RGD) cell adhesion motif just upstream its PAA (Dia and de Mejia, 2011). Peptides with an RGD motif bind integrins with high specificity, leading to antiangiogenic and anti-inflammatory effects (Kuphal et al., 2005). Cfcec^−^1 has an NGD motif at the corresponding position (Fig. 1D); as we are now set to assess the putative anticancer activity of *C. fumiferana* anionic cecropins, it will be interesting to examine the impact of mutating Asn60 to Arg in order to generate the RGD motif found in lunasin.

## Abbreviations

Cfcec^+^: *C. fumiferana* cationic cecropins
Cfcec^−^: *C. fumiferana* anionic cecropins
PAA: C-terminal poly-L-aspartic acid
AMPs: antimicrobial peptides
CAMPs: cationic antimicrobial peptides
ACPs: anticancer-peptides
APD: Antimicrobial Peptide Database
LPS: lipopolysaccharides
Pxcec^+^: cationic *Papilio Xuthus* cecropin
HccecA: *H. cecropina* cecropin-A
Pxcec^−^: anionic *P. Xuthus* cecropin
CED-9: cell-death abnormal 9
Bcl-2: B-cell lymphoma 2
APC15: subunit anaphase-promoting complex 15
MCC: the mitotic checkpoint complex
SAC: spindle assembly checkpoint
hepG-2: human liver hepatocellular carcinoma cell line.

## Acknowledgements

We thank the bioinformatics platform and the next generation sequencing platform at IBIS (Université Laval) for their assistance.

## Supplementary data

### Supplementary Material

**Figure S1.**
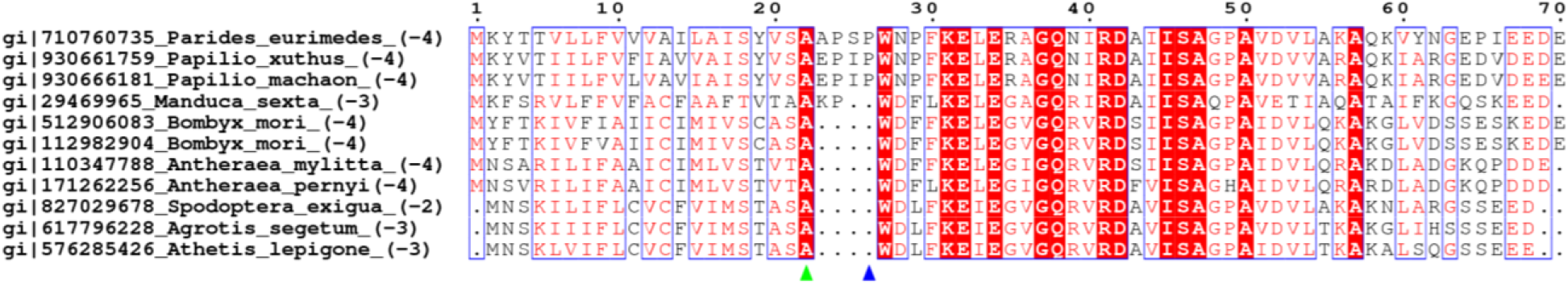
Lepidopteran cecropins with C-terminal poly-L-aspartate/glutamate. Accession number, species name and net charge (in brackets) are at the beginning of entry. Green and blue arrows identify the C-terminal amino acid of the signal peptide (SP) and the propetide (PP), respectively.

**Figure S2.**
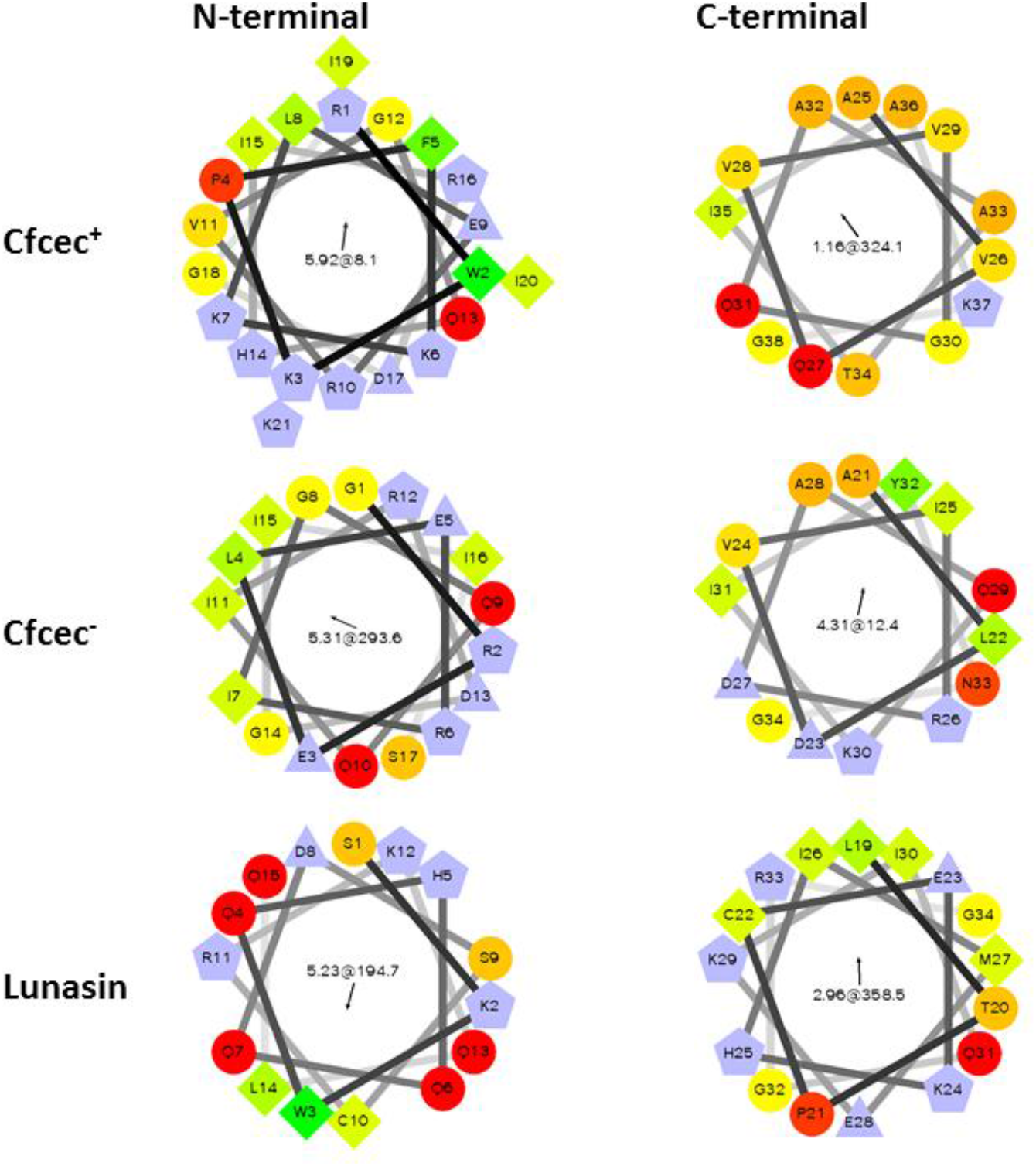
Helical-wheel diagram to illustrate the amphipathic properties of alpha helices in Cfcec^+^, Cfcec^−^ and lunasin. The plot reveals whether hydrophobic amino acids are concentrated on one side of the helix, usually with polar or hydrophilic amino acids on the other side. The hydrophobic residues as diamonds, hydrophilic residues as red circles, potentially negatively charged as triangles, and potentially positively charged as pentagons. The value in the centre of each helix represents the mean amphipathic moment <μH>.The length and the direction of the <μH> vector depend on the hydrophobicity and the position of the side chain along the helix axis. A large <μH> value means that the helix is amphipathic perpendicular to its axis. http://rzlab.ucr.edu/scripts/wheel/wheel.cgi.

**Figure S3.**
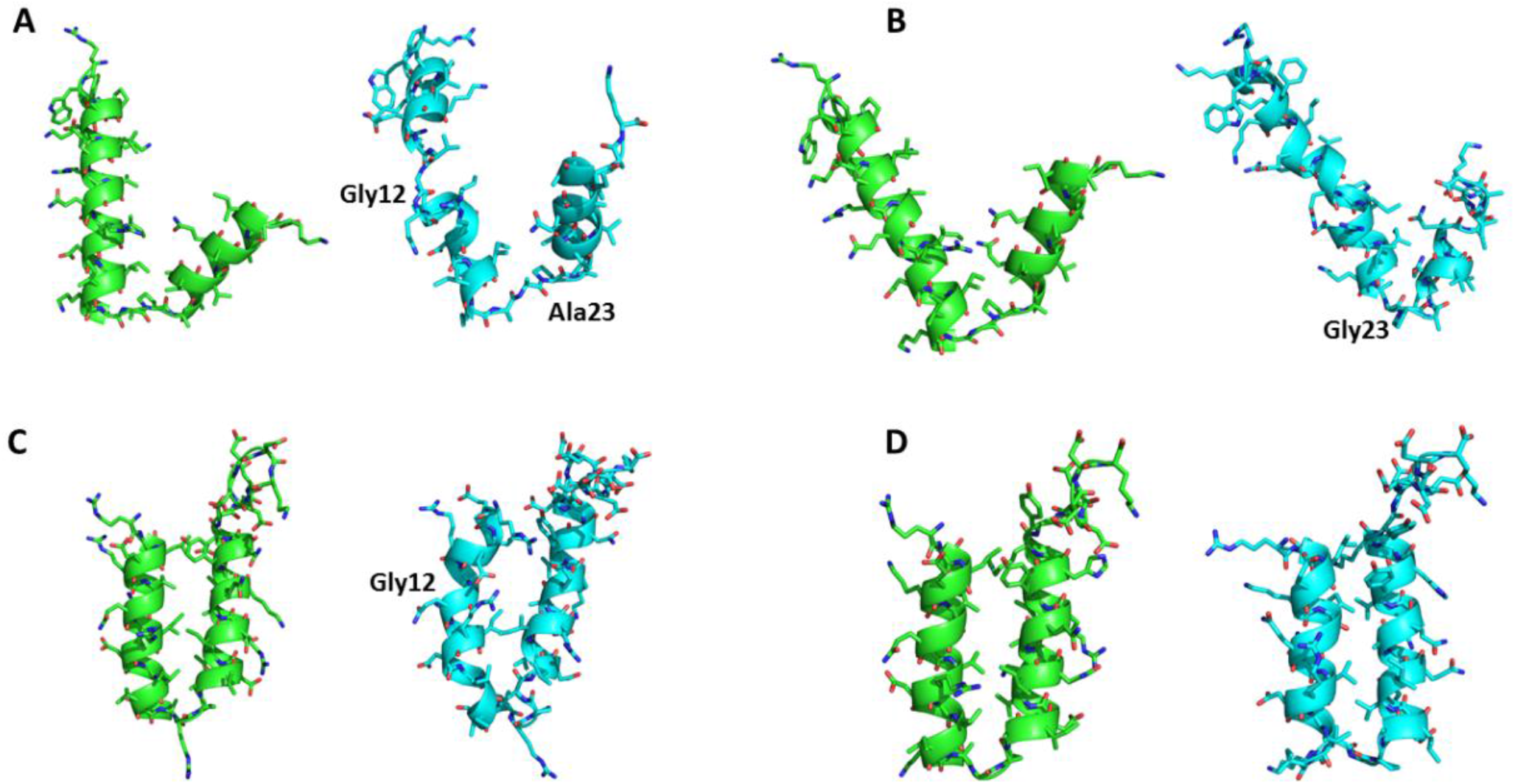
Snapshots of molecular dynamics (MD) simulation of the Cfcec^+^ and Cfcec^−^ peptides. These figures show that the flexible regions of peptides are situated at the Gly and Ala residues. (A) Cfcec^+^1, (B) Cfcec^+^2, (C) Cfcec^−^1 and (C) Cfcec^−^2. Simulation time of 0 ns (green structure) and 1 ns (cyan structure).

## References

Aerts, A.M., Carmona-Gutierrez, D., Lefevre, S., Govaert, G., François, I.E., Madeo, F.,Santos, R., Cammue, B.P., Thevissen, K., 2009. The antifungal plant defensin RsAFP2 from radish induces apoptosis in a metacaspase independent way in Candida albicans. FEBS Lett 583, 2513–2516. doi:10.1016/j.febslet.2009.07.004

Ameisen J.C., 1996. The origin of programmed cell death. Science 272, 1278–1279.

Amos S.T, Vermeer L.S, Ferguson P.M, Kozlowska J, Davy M, Bui T.T, et al. 2016. Antimicrobial Peptide Potency is Facilitated by Greater Conformational Flexibility when Binding to Gram-negative Bacterial Inner Membranes. Sci Rep. 6, 37639.

Andreu, D., Merrifield, R.B., Steiner, H., Boman, H.G., 1985. N-terminal analogues of cecropin A: synthesis, antibacterial activity, and conformational properties. Biochemistry 24, 1683–1688.

Bishop R.E., Leskiw B.K., Hodges R.S., Kay C.M., Weiner J.H., 1998. The entericidin locus of Escherichia coli and its implications for programmed bacterial cell death. J Mol Biol 280, 583–596.

Brogden, K.A., De Lucca, A.J., Bland, J., Elliott, S., 1996. Isolation of an ovine pulmonary surfactant-associated anionic peptide bactericidal for Pasteurella haemolytica. Proc. Natl. Acad. Sci. U. S. A. 93, 412–6. doi:10.1073/pnas.93.1.412

Brogden, K.A., Ackermann, M., Huttner, K.M., 1997. Small, anionic, and charge-neutralizing propeptide fragments of zymogens are antimicrobial. Antimicrob. Agents Chemother. 41, 1615–1617.

Chen, J.Y., Lin, W.J., Wu, J.L., Her, G.M., Hui, C.F., 2009. Epinecidin-1 peptide induces apoptosis which enhances antitumor effects in human leukemia U937 cells. Peptides 30, 2365–2373. doi:10.1016/j.peptides.2009.08.019

Chevreux, B., Pfisterer, T., Drescher, B., Driesel, A.J., Müller, W.E., Wetter, T., et al., 2004. Using the miraEST assembler for reliable and automated mRNA transcript assembly and SNP detection in sequenced ESTs. Genome Res14, 1147–1159.

Cho, J., Lee, D.G., 2011. The antimicrobial peptide arenicin-1 promotes generation of reactive oxygen species and induction of apoptosis. Biochim Biophys Acta 1810, 1246–1251. doi:10.1016/j.bbagen.2011.08.011

Choi, H., Lee, D.G., 2013. The influence of the N-terminal region of antimicrobial peptide pleurocidin on fungal apoptosis. J Microbiol Biotechnol 23, 1386–1394.

Dia, V.P., de Mejia, E., 2011. Lunasin induces apoptosis and modifies the expression of genes associated with extracellular matrix and cell adhesion in human metastatic colon cancer cells. Mol Nutr Food Res 55, 623–634. doi:10.1002/mnfr.201000419

Edgar, R.C., 2004. MUSCLE: a multiple sequence alignment method with reduced time and space complexity. BMC Bioinformatics 5, 113. doi:10.1186/1471-2105-5-113

Efimova, S.S., Schagina, L. V, Ostroumova, O.S., 2014. Channel-forming activity of cecropins in lipid bilayers: effect of agents modifying the membrane dipole potential. Langmuir 30, 7884–7892. doi:10.1021/la501549v

Fink, J., Boman, A., Boman, H.G., Merrifield, R.B., 1989. Design, synthesis and antibacterial activity of cecropin-like model peptides. Int J Pept Protein Res 33, 412–421.

Galvez, A.F., Chen, N., Macasieb, J., de Lumen, B.O., 2001. Chemopreventive property of a soybean peptide (lunasin) that binds to deacetylated histones and inhibits acetylation. Cancer Res 61, 7473–7478.

Galvez, A.F., de Lumen, B.O., 1999. A soybean cDNA encoding a chromatin-binding peptide inhibits mitosis of mammalian cells. Nat Biotechnol 17, 495–500. doi:10.1038/8676

Gaspar, D., Veiga, A.S., Castanho, M.A., 2013. From antimicrobial to anticancer peptides. A review. Front Microbiol 4, 294. doi:10.3389/fmicb.2013.00294

Geng, X., Harry, B.L., Zhou, Q., Skeen-Gaar, R.R., Ge, X., Lee, E.S., Mitani, S., Xue, D., 2012. Hepatitis B virus X protein targets the Bcl-2 protein CED-9 to induce intracellular Ca2+ increase and cell death in Caenorhabditis elegans. Proc Natl Acad Sci U S A 109, 18465–18470. doi:10.1073/pnas.1204652109

Hoskin, D.W., Ramamoorthy, A., 2008. Studies on anticancer activities of antimicrobial peptides. Biochim Biophys Acta 1778, 357–375. doi:10.1016/j.bbamem.2007.11.008

Hsu, J.C., Lin, L.C., Tzen, J.T., Chen, J.Y., 2011. Characteristics of the antitumor activities in tumor cells and modulation of the inflammatory response in RAW264.7 cells of a novel antimicrobial peptide, chrysophsin-1, from the red sea bream (Chrysophrys major). Peptides 32, 900–910. doi:10.1016/j.peptides.2011.02.013

Huang, Y., Feng, Q., Yan, Q., Hao, X., Chen, Y., 2015. Alpha-helical cationic anticancer peptides: a promising candidate for novel anticancer drugs. Mini Rev Med Chem 15, 73–81.

Hultmark, D., Steiner, H., Rasmuson, T., Boman, H.G., 1980. Insect immunity. Purification and properties of three inducible bactericidal proteins from hemolymph of immunized pupae of Hyalophora cecropia. Eur J Biochem 106, 7–16.

Hwang, B., Hwang, J.S., Lee, J., Kim, J.K., Kim, S.R., Kim, Y., Lee, D.G., 2011. Induction of yeast apoptosis by an antimicrobial peptide, Papiliocin. Biochem Biophys Res Commun 408, 89–93. doi:10.1016/j.bbrc.2011.03.125

Hwang, B., Hwang, J.S., Lee, J., Lee, D.G., 2011b. The antimicrobial peptide, psacotheasin induces reactive oxygen species and triggers apoptosis in Candida albicans. Biochem Biophys Res Commun 405, 267–271.

Jiang, T., Liu, M., Wu, J., Shi, Y., 2016. Structural and biochemical analysis of Bcl-2 interaction with the hepatitis B virus protein HBx. Proc Natl Acad Sci U S A 113, 2074–2079. doi:10.1073/pnas.1525616113

Jin, F., Sun, Q., Xu, X., Li, L., Gao, G., Xu, Y., Yu, X., Ren, S., 2012. CDNA cloning and characterization of the antibacterial peptide cecropin 1 from the diamondback moth, Plutella xylostella L. Protein Expr. Purif. 85, 230–238. doi:10.1016/j.pep.2012.08.006

Jin, X., Mei, H., Li, X., Ma, Y., Zeng, A.H., Wang, Y., Lu, X., Chu, F., Wu, Q., Zhu, J., 2010. Apoptosis-inducing activity of the antimicrobial peptide cecropin of Musca domestica in human hepatocellular carcinoma cell line BEL-7402 and the possible mechanism. Acta Biochim Biophys Sin 42, 259–265.

Jones, D.T., 1999. Protein secondary structure prediction based on position-specific scoring matrices. J Mol Biol 292, 195–202. doi:10.1006/jmbi.1999.3091

Kaminski G.A, Friesner R.A, Tirado-Rives J, Jorgensen W.L., 2001. Evaluation and Reparametrization of the OPLS-AA Force Field for Proteins via Comparison with Accurate Quantum Chemical Calculations on Peptides. J Phys Chem B 105, 6474–6487.

Kim, J.K., Lee, E., Shin, S., Jeong, K.W., Lee, J.Y., Bae, S.Y., Kim, S.H., Lee, J., Kim, S.R., Lee, D.G., Hwang, J.S., Kim, Y., 2011. Structure and function of papiliocin with antimicrobial and anti-inflammatory activities isolated from the swallowtail butterfly, Papilio xuthus. J Biol Chem 286, 41296–41311. doi:10.1074/jbc.M111.269225

Kuphal, S., Bauer, R., Bosserhoff, A.K., 2005. Integrin signaling in malignant melanoma. Cancer Metastasis Rev 24, 195–222. doi:10.1007/s10555-005-1572-1

Laskowski, R.A., MacArthur, M.W., Moss, D.S., Thornton, J.M., 1993. PROCHECK: a program to check the stereochemical quality of protein structures. J. Appl. Crystallogr. 26, 283–291. doi:10.1107/S0021889892009944

Lee, E., Jeong, K.W., Lee, J., Shin, A., Kim, J.K., Lee, D.G., Kim, Y., 2013. Structure-activity relationships of cecropin-like peptides and their interactions with phospholipid membrane. BMB Rep 46, 282–287.

Lee, J., Hwang, J.S., Hwang, I.S., Cho, J., Lee, E., Kim, Y., Lee, D.G., 2012. Coprisin-induced antifungal effects in Candida albicans correlate with apoptotic mechanisms. Free Radic Biol Med 52, 2302–2311. doi:10.1016/j.freeradbiomed.2012.03.012

Lee, W., Lee, D.G., 2014. Magainin 2 induces bacterial cell death showing apoptotic properties. Curr Microbiol 69, 794–801. doi:10.1007/s00284-014-0657-x

Li, L., Vorobyov, I., Allen, T.W., 2013. The different interactions of lysine and arginine side chains with lipid membranes. J Phys Chem B 117, 11906–11920. doi:10.1021/jp405418y

Li, S., Hao, L., Bao, W., Zhang, P., Su, D., Cheng, Y., Nie, L., Wang, G., Hou, F., Yang, Y., 2016. A novel short anionic antibacterial peptide isolated from the skin of Xenopus laevis with broad antibacterial activity and inhibitory activity against breast cancer cell. Arch Microbiol. doi:10.1007/s00203-016-1206-8

Lin, H.J., Huang, T.C., Muthusamy, S., Lee, J.F., Duann, Y.F., Lin, C.H., 2012. Piscidin-1, an antimicrobial peptide from fish (hybrid striped bass morone saxatilis x M. chrysops), induces apoptotic and necrotic activity in HT1080 cells. Zool. Sci 29, 327–332. doi:10.2108/zsj.29.327

Lu, Y., Zhang, T.F., Shi, Y., Zhou, H.W., Chen, Q., Wei, B.Y., Wang, X., Yang, T.X., Chinn, Y.E., Kang, J., Fu, C.Y., 2016. PFR peptide, one of the antimicrobial peptides identified from the derivatives of lactoferrin, induces necrosis in leukemia cells. Sci Rep 6, 20823. doi:10.1038/srep20823

Ma, N.F., Lau, S.H., Hu, L., Xie, D., Wu, J., Yang, J., Wang, Y., Wu, M.C., Fung, J., Bai, X., Tzang, C.H., Fu, L., Yang, M., Su, Y.A., Guan, X.Y., 2008. COOH-terminal truncated HBV X protein plays key role in hepatocarcinogenesis. Clin Cancer Res 14, 5061–5068. doi:10.1158/1078-0432.CCR-07-5082

Mansfeld, J., Collin, P., Collins, M.O., Choudhary, J.S., Pines, J., 2011. APC15 drives the turnover of MCC-CDC20 to make the spindle assembly checkpoint responsive to kinetochore attachment. Nat Cell Biol 13, 1234–1243. doi:10.1038/ncb2347

Meng, M.X., Ning, J.F., Yu, J.Y., Chen, D.D., Meng, X.L., Xu, J.P., Zhang, J., 2014. Antitumor activity of recombinant antimicrobial peptide penaeidin-2 against kidney cancer cells. J Huazhong Univ Sci Technol. Med Sci 34, 529–534. doi:10.1007/s11596-014-1310-4

Oh, D., Shin, S.Y., Lee, S., Kang, J.H., Kim, S.D., Ryu, P.D., Hahm, K.S., Kim, Y., 2000. Role of the hinge region and the tryptophan residue in the synthetic antimicrobial peptides, cecropin A(1-8)-magainin 2(1-12) and its analogues, on their antibiotic activities and structures. Biochemistry 39, 11855–11864.

Otvos, L., 2000. Antibacterial peptides isolated from insects. J Pept Sci 6, 497–511. doi:10.1002/1099-1387(200010)6:10<497::AID-PSC277>3.0.CO;2-W

Park, C., Lee, D.G., 2010. Melittin induces apoptotic features in Candida albicans. Biochem Biophys Res Commun 394, 170–172.

Ren, J., Wen, L., Gao, X., Jin, C., Xue, Y., Yao, X., 2009. DOG 1.0: illustrator of protein domain structures. Cell Res 19, 271–273. doi:10.1038/cr.2009.6

Ren, S.X., Cheng, A.S., To, K.F., Tong, J.H., Li, M.S., Shen, J., Wong, C.C., Zhang, L., Chan, R.L., Wang, X.J., Ng, S.S., Chiu, L.C., Marquez, V.E., Gallo, R.L., Chan, F.K., Yu, J., Sung, J.J., Wu, W.K., Cho, C.H., 2012. Host immune defense peptide LL-37 activates caspase-independent apoptosis and suppresses colon cancer. Cancer Res 72, 6512–6523. doi:10.1158/0008-5472.CAN-12-2359

Riedl, S., Zweytick, D., Lohner, K., 2011. Membrane-active host defense peptides--challenges and perspectives for the development of novel anticancer drugs. Chem Phys Lipids 164, 766–781. doi:10.1016/j.chemphyslip.2011.09.004

Saitou, N., Nei, M., 1987. The neighbor-joining method: a new method for reconstructing phylogenetic trees. Mol Biol Evol 4, 406–425.

Steiner, H., Hultmark, D., Engström, A., Bennich, H., Boman, H.G., 1981. Sequence and specificity of two antibacterial proteins involved in insect immunity. Nature 292, 246–248.

Suttmann, H., Retz, M., Paulsen, F., Harder, J., Zwergel, U., Kamradt, J., Wullich, B., Unteregger, G., Stöckle, M., Lehmann, J., 2008. Antimicrobial peptides of the Cecropin-family show potent antitumor activity against bladder cancer cells. BMC Urol 8, 5. doi:10.1186/1471-2490-8-5

Tajima, F., Nei, M., 1984. Estimation of evolutionary distance between nucleotide sequences. Mol Biol Evol 1, 269–285.

Tamura, K., Stecher, G., Peterson, D., Filipski, A., Kumar, S., 2013. MEGA6: Molecular Evolutionary Genetics Analysis version 6.0. Mol Biol Evol 30, 2725–2729. doi:10.1093/molbev/mst197

Wang, Y., Qu, L., Gong, L., Sun, L., Gong, R., Si, J., 2013. Targeting and eradicating hepatic cancer cells with a cancer-specific vector carrying the Buforin II gene. Cancer Biother Radiopharm 28, 623–630. doi:10.1089/cbr.2012.1469

Webb, B., Sali, A., 2014. Comparative Protein Structure Modeling Using MODELLER. Curr Protoc Bioinforma. 47, 5.6.1–32. doi:10.1002/0471250953.bi0506s47

White, S.H., Wimley, W.C., 1998. Hydrophobic interactions of peptides with membrane interfaces. Biochim Biophys Acta 1376, 339–352.

Wu, C., Geng, X., Wan, S., Hou, H., Yu, F., Jia, B., Wang, L., 2015. Cecropin-P17, an analog of Cecropin B, inhibits human hepatocellular carcinoma cell HepG-2 proliferation via regulation of ROS, Caspase, Bax, and Bcl-2. J Pept Sci 21, 661–668. doi:10.1002/psc.2786

Xu, D., Jaroszewski, L., Li, Z., Godzik, A., 2014. FFAS-3D: improving fold recognition by including optimized structural features and template re-ranking. Bioinformatics 30, 660–667. doi:10.1093/bioinformatics/btt578

Yi, H.Y., Chowdhury, M., Huang, Y.D., Yu, X.Q., 2014. Insect antimicrobial peptides and their applications. Appl Microbiol Biotechnol 98, 5807–5822. doi:10.1007/s00253-014-5792-6

Zasloff, M., 2002. Antimicrobial peptides of multicellular organisms. Nature 415, 389–395. doi:10.1038/415389a

Zhang, H.T., Wu, J., Zhang, H.F., Zhu, Q.F., 2006. Efflux of potassium ion is an important reason of HL-60 cells apoptosis induced by tachyplesin. Acta Pharmacol Sin 27, 1367–1374. doi:10.1111/j.1745-7254.2006.00377.x

